# The nematode symbiotic bacterium *Xenorhabdus griffiniae* can sense and respond to the presence of its host *Steinernema hermaphroditum*

**DOI:** 10.1101/2025.06.16.660008

**Authors:** Elin M. Larsson, Carly R. Myers, Dianne K. Newman, Mengyi Cao

**Affiliations:** Division of Biology and Bioengineering, California Institute of Technology, Pasadena, CA, USA; Division of Biosphere Sciences and Engineering, Carnegie Institution for Science, Pasadena, CA, USA; Division of Geological and Planetary Sciences, California Institute of Technology, Pasadena, CA, USA

**Keywords:** host-microbe interactions, signaling, RNA-sequencing, ymdA

## Abstract

The mutualistic symbiosis between *Xenorhabdus* bacteria and *Steinernema* nematodes has both ecological and practical importance. Their relationship co-evolved to make them dependent on each other in their life cycle where they kill, feed and reproduce within insect larvae together. This behavior has shed light on them as promising candidates to replace conventional pesticides. Despite the importance of the *Xenorhabdus* in insect-killing, the mechanisms by which the bacteria might sense and respond to the presence of their nematode host are not well understood. We performed an RNA-sequencing experiment on the bacterial partner, *X. griffiniae*, in close proximity, but not in direct contact with their host nematode, *Steinernema hermaphroditum*, to identify differentially regulated genes in these conditions, followed by genetic analysis to determine their functional significance. We show that *X. griffiniae* changes its transcriptomic profile in a small number of genes in the presence of axenic nematodes, but not in the presence of nematodes that are already colonized by *X. griffiniae*. We select the most differentially regulated gene, *ymdA*, for further investigation, and show that it plays a role in biofilm formation and affects host colonization efficiency. This work advances our understanding of bacterial sensing of nematodes and motivates future research in deepening our understanding of this underexplored ecological interaction.

**Importance:** Interactions between different organisms in the soil environment are enabled by the ability to sense and respond to organisms nearby. In many cases, the underlying sensing mechanisms that mediate these ecologically important interactions are not known. Filling in the knowledge gap of how these interactions occur can help us understand how microbial populations organize themselves, establish symbiotic relationships, or compete with other species in complex environments. Furthermore, elucidating these mechanisms may inform the development of new tools in agriculture, such as improved biological control agents. Here we use the bacterial symbiont *Xenorhabdus griffiniae* of the entomopathogenic nematode *Steinernema hermaphroditum* to study whether symbiotic bacteria of nematodes are able to sense their presence. Our results suggest that *X. griffiniae* can sense the presence of its nematode host and respond by modulating the gene expression level of several genes, where at least one, *ymdA*, has implications for host colonization.

## Introduction

Nematodes are the most ubiquitous metazoans on earth and play key roles in diverse ecosystems, through nutrient cycling, predation, pathogenesis, and parasitism with other types of organisms [1, 2, 3, 4]. Nematodes’ ability to sense and be sensed by other organisms enable many of these ecosystem interactions. Some examples include the ability of *Caenorhabditis elegans* to sense and respond to its bacterial prey [5], and bacterial pathogens [6, 7], and the ability of plants to trigger an immune response in the presence of plant-pathogenic nematodes [8].

Many of these ecosystem interactions occur between nematodes and bacteria. Although the ability of nematodes to sense bacteria is well documented [5, 6, 7], much less is understood about how bacteria sense nematodes. Some bacteria are able to import and metabolize nematode signaling molecules called ascarosides [9], but to our knowledge, the only documented example of bacterial *sensing* of a nematode is for the nematode-predatory bacterium *Bacillus nematocida*. This bacterium initiates production of volatile attractants and sporulation upon sensing the *C. elegans*-made gas morpholine in order to germinate and feed on the nematode from within, after surviving engulfment [10]. It is therefore an open question whether bacteria can sense nematodes in non-predation contexts, such as mutualistic relationships.

Entomopathogenic (insect-parasitic) nematodes of the genus *Steinernema* are distinguished by having a core microbiome dominated by a single bacterial species, with strong specificity between each nematode species and its preferred symbiont [11, 12]. During the infective juvenile (IJ) stage, the symbiotic bacterium resides inside an intestinal pocket termed the receptacle, while the nematode seeks and infects insect prey. Once inside the insect, *Xenorhabdus* bacteria are released from the IJs and contribute to insect killing. In the insect cadaver, the bacterial symbionts play a major role in supporting nematode development [13, 14, 15], providing nutrients and producing antimicrobial molecules that aid in protecting the cadaver from the surrounding soil microbiome [16, 17, 18]. After depleting the nutrients in the insect cadaver, symbionts reassociate with nematode progeny in the IJ stage, and leave the insect cadaver to seek a new insect host to infect.

The most well studied pair is *Steinernema carpocapsae* and *Xenorhabdus nematophila*, where the ability for the nematodes to recognize their bacterial symbiotic partners, through the bacterial expression of the *nilABC* genes, has been described [19]. To our knowledge there has been no investigation, in these or other *SteinernemaXenorhabdus* pairs, of whether the bacterial symbionts can sense and respond to the presence of their nematode hosts and if so how. In this work, we leverage the emerging genetic model entomopathogenic nematode symbiont pair *Steinernema hermaphroditum* and *Xenorhabdus griffiniae* HGB2511 to determine whether *X. griffiniae* can sense and respond to the presence of its nematode host by recognizing nematode secreted molecules.

## Methods

### Preparation of nematodes for trans-well experiment

*Steinernema hermaphroditum* nematodes were maintained as described in Cao et al. [20]. Briefly, WT (PS9179) conventional infective juveniles (IJs) were produced by natural propagation of *S. hermaphroditum* through 5th instar larvae of *Galleria mellonella* (Grubco, OH) and collected by a modified White Trap [21]. WT and *daf22* mutant (MCN0006) *S. hermaphroditum* were cryopreserved at -80^*°*^C and passed on Nematode Growth Media (NGM) agar plates [22] and then used for preparing axenic IJs. To produce axenic (gnobiotic) IJs, nematodes were grown on the lawns of *X. griffiniae* HGB2511 on liver kidney agar (50 g beef liver, 50 g beef kidney blended in 50 ml water, 2.5 g NaCl, 7.5 g agar: modified from [13]). Gravid adult hermaphrodites were harvested to extract axenic eggs using a modified protocol from (cite), which were then seeded on beef liver kidney agar. Axenic IJs were trapped and stored in water at less than 8 IJs per *µ*l until being used for trans-well experiments.

To optimize trans-well experimental conditions, high density of IJ nematode pellets were tested in sterile water and various media including nematode growth media (NGM), M9 (with and without glucose or cholesterol supplementation), and at various temperatures of 22^*°*^C, 25^*°*^C, and 30^*°*^C. M9 glucose supplemented with 0.1% cholesterol at 25^*°*^C yielded optimal IJ viability after overnight incubation in the trans-wells and was a feasible condition for the growth of *Xenorhabdus* bacteria, therefore was used for the trans-well experiment. Before starting the experiment, freshly emerged (less than two weeks of aging) conventional and axenic IJs were surface sterilized using 0.5% bleach for 90 seconds to remove the outer cuticle which facilitates the releasing of secreted molecules. The bleaching treatment was followed by three washes in water, and IJs were stored in water at less than 5 IJs per *µ*l overnight at 22^*°*^C. Right before the experiment, IJs were washed with M9 medium with glucose (Teknova) supplemented with 0.1% cholesterol, then concentrated by centrifugation, and transferred into trans-wells at high density (approximately 20006000 IJs per trans-well). Conventional IJs were propagated and emerged from three batches of insects and were used as three independent biological replicates. Axenic IJs prepared from three independent batches of nematode stocks through three rounds of axenic IJ extraction were used as three biological replicates. The density and mortality of IJ nematodes were assessed by microscopy (Supplemental Table S4).

### Trans-well experiment

Starting from three individual colonies, three bacterial overnight cultures were inoculated into 5 ml of LB supplemented with 1% sodium pyruvate and grown in a shaker at 30^*°*^C. The following morning, the bacterial cultures were washed twice in M9 medium with glucose (Teknova) supplemented with 0.1% cholesterol. The cultures were then diluted to a final OD of 0.05 in a volume of 1.2 ml and transferred to a 24 well plate (VWR). A 0.4 *µ*m membrane insert was added to each well and 50 *µ*l of dense nematode pellet was added to each membrane insert, as well as 50 *µ*l of M9 cholesterol medium. In the “no nematode condition” the membrane inserts were left empty. The bacterial cultures were grown shaking at 100 rpm for 24 hours at 25^*°*^C. The bacteria were then pelleted at 8000 rpm and flash frozen using liquid nitrogen and stored at -80C. Bacterial pellets were sent to the microbial sequencing and analysis center at the University of Pittsburgh for RNA extraction and sequencing. After the experiment, IJs were transferred from the trans-well inserts to recover in water overnight.

### Data analysis and selection of target genes

The RNA sequences were mapped to the X. griffiniae HGB2511 reference genome [23] that was annotated using Prokka 1.14.6 [24]. The abundance quantification of the transcripts was done using kallisto [25], with the default settings except for the boot strap samples that were set to 100. To identify differentially expressed genes, DESeq2 was used [26], using the default settings for calculating the adjusted p-values. We chose a cutoff at an adjusted p-value of 0.05 and fold change of 2 to select genes of interest. The full data set is publicly available through the NCBI gene expression omnibus (GEO GSE299917).

### Construction of deletion mutants, overexpression, and fluorescently labelled strains

Deletion strains were constructed by replacing the gene of interest with a kanamycin resistance cassette using allelic replacement. Briefly, 1500 base pairs upstream and downstream of the gene were PCR amplified and cloned into the pKR100 plasmid, sandwiching the kanamycin cassette. A kanamycin cassette was placed in between the homology sequences and inserted into the genome, replacing the deleted gene. To make plasmids used for genome insertions, standard 3G cloning was used [27].

Both deletion and insertion plasmids were transferred to *X. griffiniae* by conjugation. Starting from a single colony, *Xenorhabdus griffiniae, E. coli* helper (only used for insertions) and *E. coli* donor were grown overnight in 5 ml LB broth (supplemented with antibiotics and 0.3 mM diaminopimelic acid, DAP, as needed). The optical density (OD600) of each strain was measured. The volume to obtain OD600 3 for each strain was calculated and pelleted. The pellets were then washed twice with LB to remove residual antibiotics. The pellets were combined by sequentially resuspending them in 100 *µ*l of LB until all pellets were mixed. The cells were then pipetted onto filter membrane (MF-Millipore) on an agar plate and incubated at 30^*°*^C overnight. The following day, the filter was transferred to 5 ml of LB in a conical tube (Falcon) and vortexed until the bacteria on the filter paper were resuspended in the liquid. The liquid was then plated in serial dilutions onto LB agar plates supplemented with kanamycin (50 *µ*g/ml) or streptomycin (25 *µ*g/ml) and incubated in 30^*°*^C overnight. Colonies of *X. griffiniae* transconjugants were picked and verified by colony PCR and sequencing.

### Crystal violet biofilm assay

The crystal violet assay is modified from a microtiter biofilm protocol [28]. Starting from an overnight LB culture, bacterial strains were diluted in LB to OD600 0.003. 1 ml of diluted culture was added to a 6 ml borosilicate glass tube and incubated statically for 4 days at room temperature (RT). The cultures were then decanted and the optical density of the liquid was measured. The tubes were washed twice by submerging them in MilliQ water and tapped on a paper towel to remove most of the residual water. After washing, 1.2 ml of crystal violet (0.1% w/v) was added to each tube. The tubes were incubated for 15 minutes at RT and rolled gently at a slight angle twice during the incubation. The crystal violet was then removed by decanting and the tubes were again washed twice by submerging in MilliQ water. The tubes were then tapped against a paper towel and dried for 24 hours at RT. After drying, 1.2 ml of acetic acid (30% v/v) was added to each tube. The tubes were incubated for 10 minutes at RT with gentle rolling of the tubes twice during the incubation. After incubation, the dissolved crystal violet/acetic acid mixture was measured using a Biotek plate reader at 595 nm.

### Attachment assay - microcolony formation

Overnight LB cultures of WT, ∆*ymdA* and the *ymdA* overexpression strain were diluted to OD600 0.01 into M9 medium with glucose (Teknova). 400 *µ*l culture was transferred to an 8 well glass bottom plate and incubated at RT for 20 hours. The bottom of each well was then imaged using a Nikon Ti2 microscope at 40x magnification.

### Nematode colonization

The nematode colonization protocol was modified from St. Thomas et al. [29].

#### Preparation of bacteria

Bacterial strains for the colonization experiment were streaked from glycerol stocks onto agar plates containing appropriate antibiotics and incubated at 30^*°*^C. The following day, a colony was used to inoculate 5 ml of LB supplemented with antibiotics and grown for 16 hours. The optical density of the cultures was then measured. The competing strains in each condition were mixed at a 1:1 ratio to end up with equal ODs for each strain. 700 *µ*l of the strain mix was added to a liver kidney plate with added kanamycin (50 *µ*g/ml). The plates were then incubated at 30^*°*^C overnight before adding nematodes.

#### Nematode colonization and collection

The nematode colonization protocol was modified from Murfin et al. [30]. 2 *µ*l of surface sterilized (1% bleach) nematodes were added to a microscope slide in triplicate. The number of nematodes was then counted and the concentration (nematode/*µ*l) in the stock solution was calculated.

A volume corresponding to 200 IJs was added to each bacterial lawn on the liver kidney agar plates with bacterial lawns, which were then incubated at 25^*°*^C for 7 days. The nematodes were then water trapped by placing the 60 mm liver kidney agar plates in a 100 mm x 200 mm deep petri dish where the bottom was covered in autoclaved water. After 7 additional days, nematodes trapped in the water could be collected for further experiments.

### CFU plating from nematode grounds

After surface sterilizing 200 IJs, the nematodes were ground up in a total volume of 200 *µ*l of LB by bead (stainless steel, 6 mm diameter) beating for 6 minutes at 50 s^*−*1^ amplitude (Qiagen, TissueLyser bead beater). The grounds were then serially diluted and plated on selective agar plates and incubated in room temperature for two days before colony counting.

### Fluorescence microscopy of whole nematodes

Nematodes were collected from the water traps, then they were concentrated to a volume of 300 *µ*l. To this, 5 *µ*l of levamisole (200 *µ*M) was added to the nematodes to paralyze them. The nematodes were imaged using a Nikon Ti2 microscope at 10x magnification, using phase contrast, GFP, and RFP channels. At least 100 nematodes were imaged per condition and replicate (5 technical replicates per biological replicate, where each technical replicate is a nematode population from a separate liver kidney agar plate).

## Results

### *X. griffiniae* exhibits a transcriptional response to axenic *Steinernema hermaphroditum* infective juveniles

To determine whether *X. griffiniae* can sense the presence of its nematode host, we focused on the IJ stage of *S. hermaphroditum* as the source of chemical signals to be recognized because of its biological relevance and experimental feasibility. In the natural life cycle of *Steinernema* within an insect cadaver, nematodes are usually mixed in their developmental stages. The bacterial symbiont does not have a free living stage, but is outside of the nematode during two parts of the life cycle: the insect colonization and nutrient depletion phases (Figure 1A). The ability to sense the nematode host in the latter might be important because successful colonization of the nematode during this phase allows the bacterium to disperse out into the wider soil environment. We therefore hypothesized that *X. griffiniae* can sense the presence of *Steinernema hermaphroditum*, specifically by sensing chemicals secreted by *Steinernema hermaphroditum* in the IJ stage.

**Figure 1:**
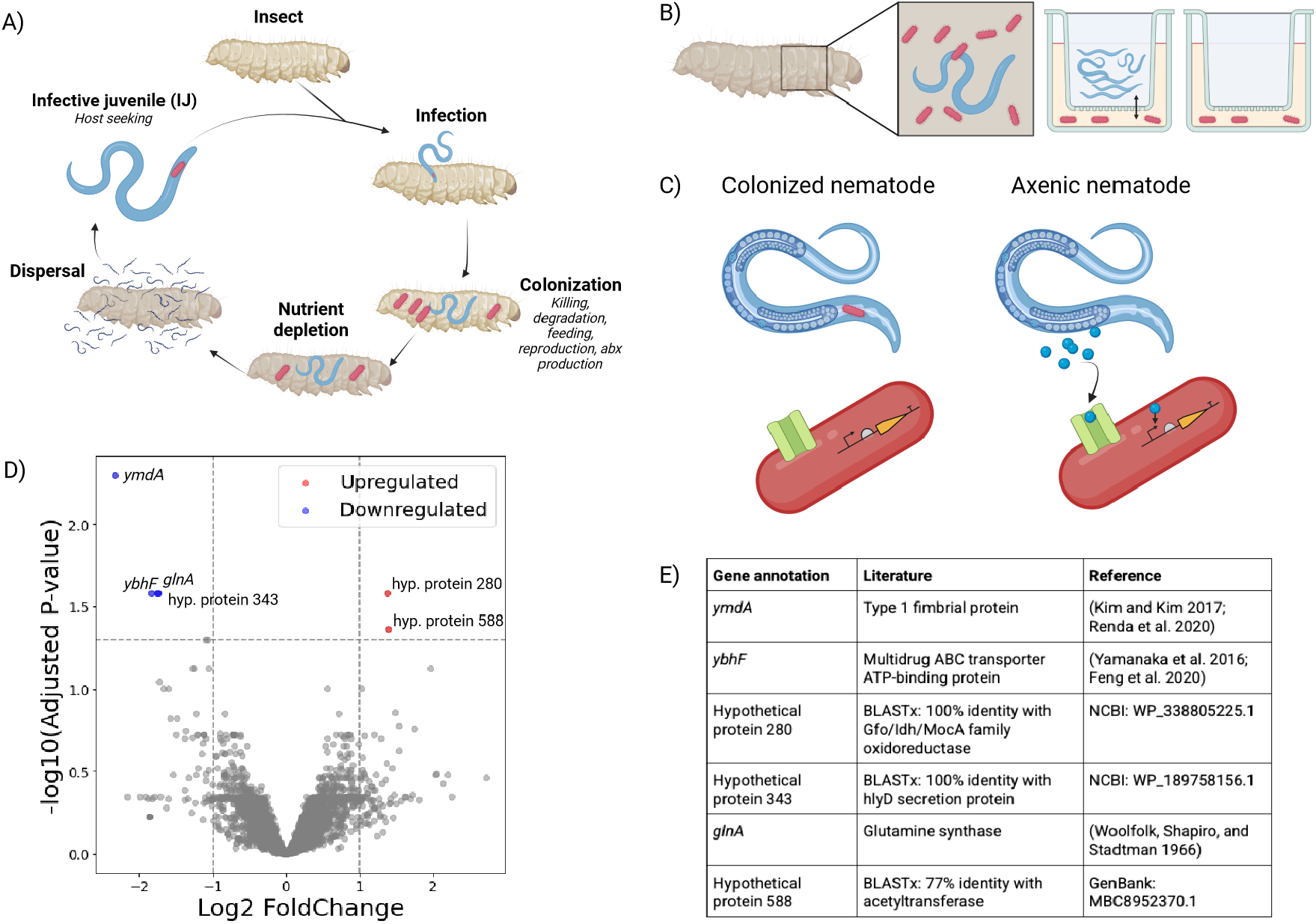
(A) *Steinernema-Xenorhabdus* life-cycle overview. (B) The trans-well experiment co-cultures *X. griffiniae* with IJ-stage *S. hermaphroditum* in order to match the life stage in which the bacteria and nematodes would naturally be apart from each other. A trans-well set-up allows cross-feeding of chemicals, but prevents direct contact between nematodes and bacteria. (C) Schematic of proposed model for signal secretion and sensing. In this model, only axenic nematodes secrete signaling molecules that can be sensed by bacteria. (D) Volcano plot of differential expression in *Xenorhabdus* co-cultured with axenic wildtype *S. hermaphroditum*.(E) Table listing differentially expressed genes in *X. griffiniae* with axenic WT nematodes. nematodes are present. The gene for a third hypothetical protein along with *ymdA, ybhF*, and *glnA* are downregulated in the presence of axenic nematodes. These results suggest that *X. griffiniae* can indeed sense the presence of its nematode host *S. hermaphroditum*, but it can only do so when the host is not colonized by bacteria.

To test this hypothesis, we developed a trans-well assay where the nematodes are separated from the bacteria by a membrane, preventing engulfment, that allows diffusion of potential chemical signals involved in sensing (Figure 1B). This allows us to isolate the sensing of nematode secreted molecules from factors like physical contact or nutrients contained in specific nematode tissues.

We compared pure cultures of *X. griffiniae* against co-cultures with colonized or axenic *Steinernema hermaphroditum* in the trans-well insert, in two genetic backgrounds for the nematode (wildtype and *daf-22* deficient [31]). This mutant was included because the *daf-22* gene is involved in the biosynthetic pathway for ascarosides. Since some bacteria are known to metabolize these molecules, we hypothesized that this class of molecules could potentially be sensed by bacteria and that including a nematode lacking the natural ascaroside composition might provide additional information.

After co-culturing the bacteria and nematodes, we sequenced the bacterial RNA to determine whether the presence of the nematodes altered the transcriptome of the bacteria. We found that *X. griffiniae* only shifted its gene expression profile in response to axenic nematodes, and not to colonized nematodes (Figure 1C and Figure S1-S2). Furthermore, only a small number of genes were differentially expressed in the presence of the axenic wildtype nematodes (Figure 1D-E, Table S3 for axenic *daf-22* dataset). Genes for two hypothetical proteins are upregulated when axenic

To better understand the nature of *X. griffiniae*’s response to the presence of *Steinernema hermaphroditum*, we chose the most significantly differentially expressed gene from our dataset, a putative homologue of *ymdA*, for further characterization (Figure 1D-E).

### Overexpressing *ymdA* leads to a biofilm defect *in vitro*

In *Escherichia coli, ymdA* is a type 1 fimbrial protein that is known to be involved in biofilm formation [32, 33], but its role in *X. griffiniae* has not been studied. Biofilm formation plays a key role in attachment, survival, and persistence in the host animal for both pathogenic and mutualistic host-microbe associations. To determine if *ymdA* similarly plays a role in biofilm formation in *X. griffiniae*, we performed an *in vitro* biofilm formation experiment with WT, *ymdA* deletion, and *ymdA* overexpression strains. Quantification of crystal violet staining of the biofilm shows that deleting *ymdA* does not significantly affect biofilm formation *in vitro*, but overexpressing *ymdA* significantly reduces the cells’ ability to form stable biofilms (Figure 2A-B). This effect is likely not due to growth a burden associated with *ymdA* overexpression, as growth curves indicate that the *ymdA* overexpression strain grows to a similar final density as WT in static liquid cultures (Figure 2C).

**Figure 2:**
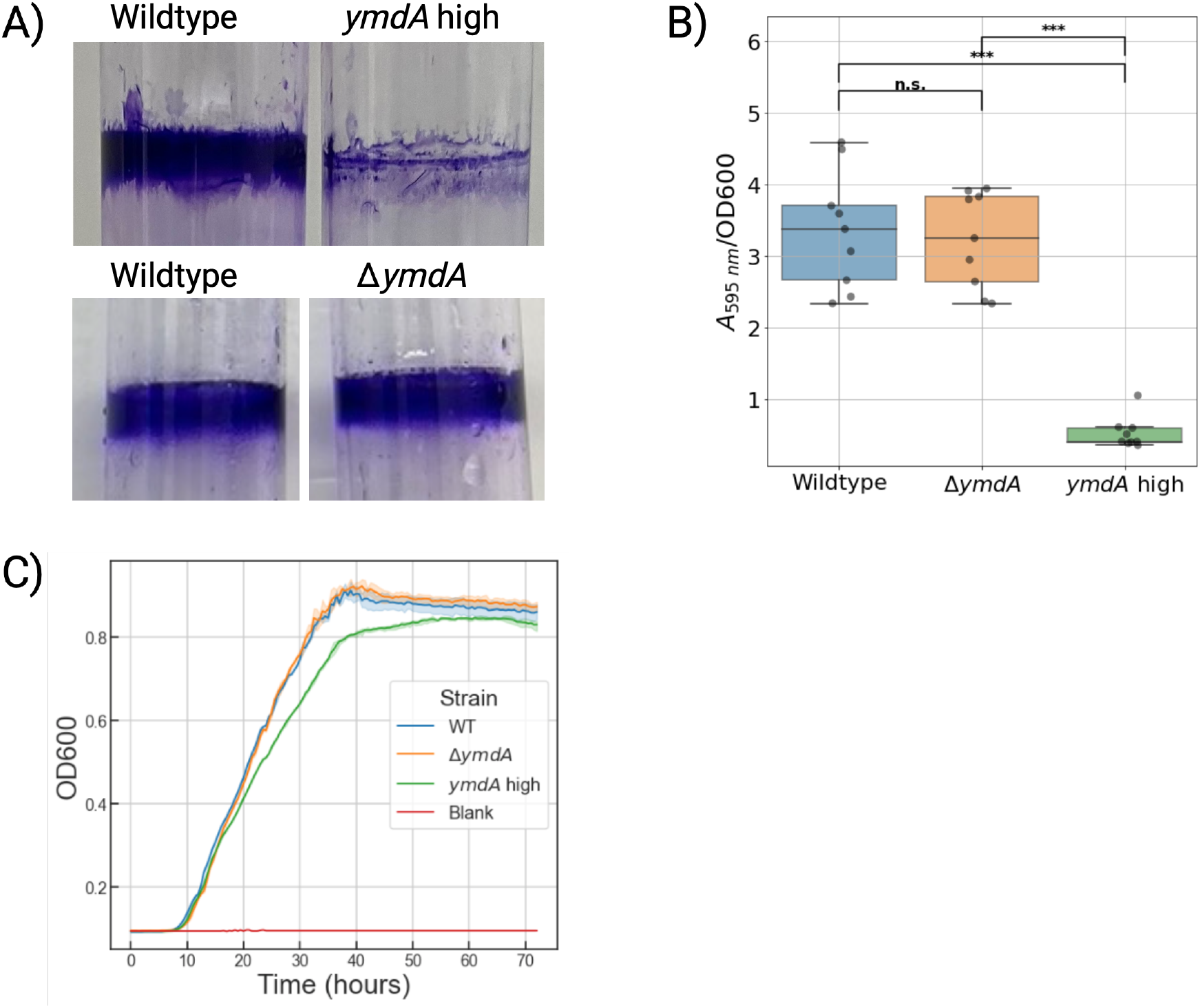
(A) Representative image of crystal violet stained biofilms stained after 3 days of static incubation (upper left: WT, upper right: *ymdA* overexpression strain, lower left WT, lower right ∆*ymdA*. (B) Destained biofilms are quantified by measuring crystal violet absorbance at 595 nm normalized by measured optical density. Technical triplicates from three biological replicates are plotted. Welch ttest p-value not significant between WT and ∆*ymdA* and <0.001 for WT and *ymdA* overexpression. (C) Growth curves for static growth in LB.

The RNA-sequencing experiment shows that, *ymdA* is downregulated in the presence of the nematodes. Accordingly, we hypothesized that *X. griffiniae* senses and responds to axenic IJs by suppressing *ymdA* expression, thereby modulating cell-cell interactions that are disadvantageous in the establishment of symbiosis. To test this hypothesis, we began by attempting to pinpoint the stage in biofilm development where YmdA acts.

### The *ymdA* overexpression strain forms large microcolonies in early attachment

Several studies show that an increase in fimbrial expression leads to enhanced biofilm formation [34, 35, 36, 37]. However, for *ymdA* specifically, previous studies showed that overexpression leads to a biofilm defect in *E. coli* [32, 33]. Our crystal violet experiment suggested that *ymdA* might function similarly in *X. griffiniae* as in *E. coli*.

Deficiencies in biofilm formation can occur via a number of mechanisms at different stages of development, such as inhibiting early attachment or blocking the secretion of exopolymers required for stable macroscale biofilm growth. To begin, we performed a microscopy assay to assess the impact of *ymdA* on the ability of *X. griffiniae* to form microcolonies, an early attachment stage in biofilm development.

Interestingly, the *ymdA* overexpression strain formed larger and more clustered microcolonies than WT and *ymdA* deletion strains (Figure 3), suggesting that *ymdA* expression can indeed promote cell-cell adherence similarly to bacteria increasing expression of other fimbrial proteins. WT and the deletion mutant microcolony patterns were similar, which might mean that *ymdA* is not required for microcolony formation, or that other fimbrial proteins annotated on the *X. griffiniae* genome (*mrpA, yfcS, yfcP, lpfB, lpfA*1/2 and *mrkD*) might be expressed to compensate for the lack of *ymdA* in the deletion strain. These results together suggest that in *X. griffiniae*, high levels of *ymdA* promote cell-cell adherence early at the microcolony scale, but inhibit macroscale biofilm formation that occurs over several days. But how is this relevant to a symbiotic life cycle? To answer this question, we turned to a nematode colonization assay.

**Figure 3:**
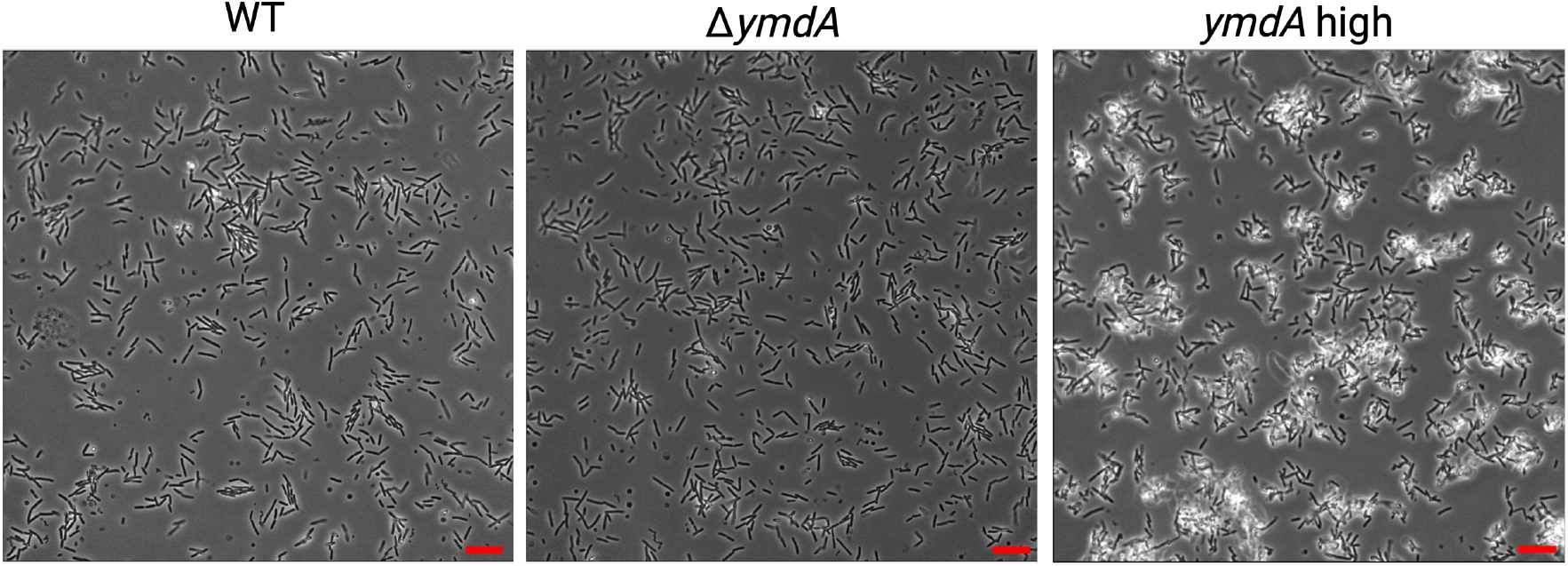
Representative images of WT, deletion mutant and overexpression strain cells at 20 hours of growth. Scale bar 5 *µ*m.

### *ymdA* overexpression results in a disadvantage in competitive nematode colonization

While there are multiple points where biofilm development could impact *Xenorhabdus*-*Steinernema* symbiosis, we reasoned that the first step of the interaction—colonization of the host—was one that might be particularly sensitive to bacterial cell-cell interactions. To determine if *ymdA* in *X. griffiniae* affects host association, we performed a competitive colonization assay where WT bacteria were mixed at equal cell density with the *ymdA* overexpression strain (Figure 4A). A competitive colonization assay provides an opportunity to reveal subtle phenotypes that individual colonization of WT and mutants may not show. Additionally, the competitive set-up better mimics the setting in the nutrient depleted insect cadaver, where only a fraction of the bacteria in the cadaver will be able to re-colonize the nematode host. The question at hand is what bacteria in a large population of bacteria are able to most efficiently and robustly colonize the nematodes.

**Figure 4:**
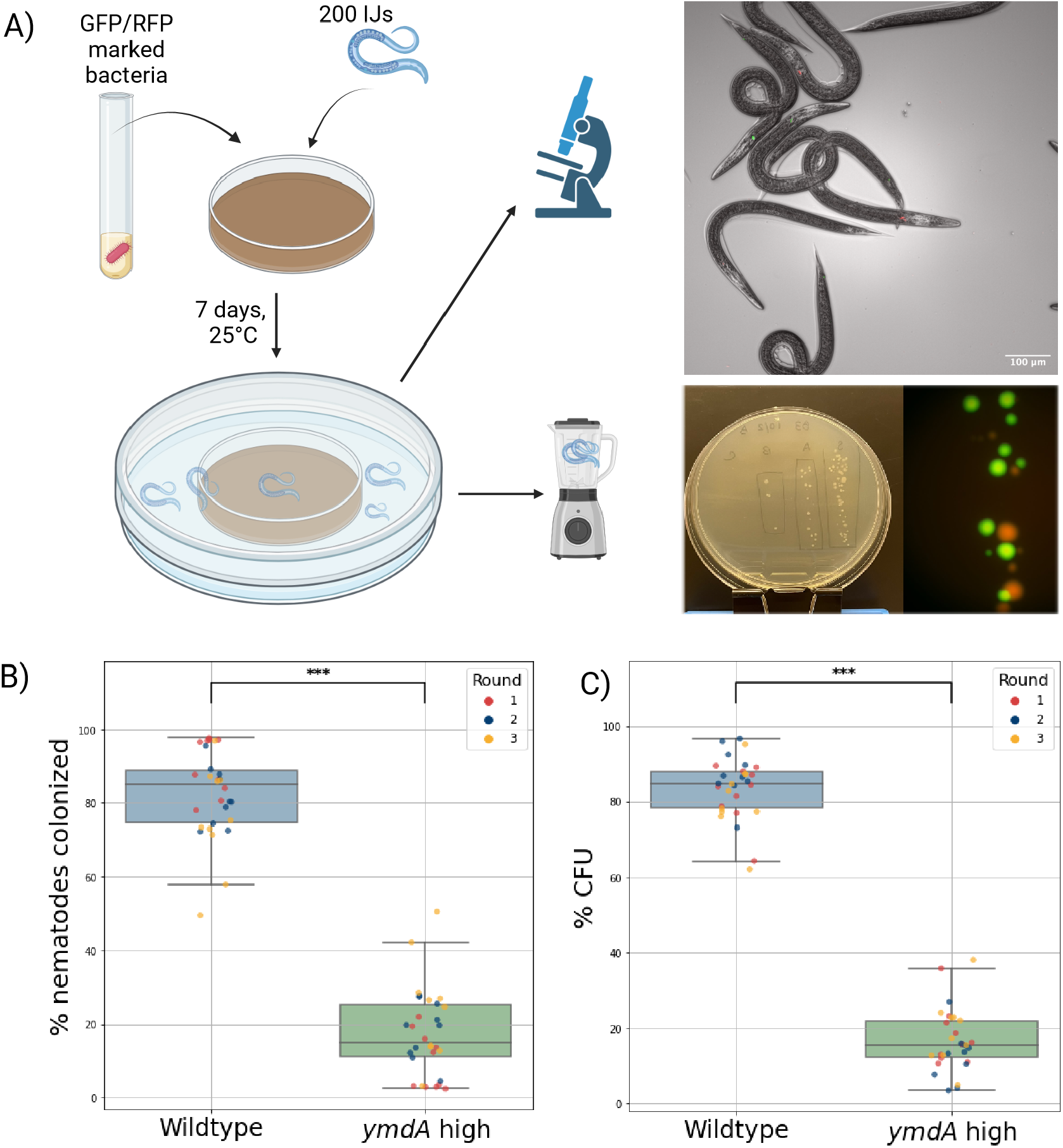
(A) Schematic of the experimental procedure for the colonization experiments and representative images of microscopy and colony imaging. (B) Percentages of nematodes colonized by WT and *ymdA* overexpression strain bacteria. Ten replicates per biological replicate (*n* = 3) are plotted. (C) Percentages of CFUs from WT and *ymdA* overexpression strain bacteria. Ten replicates per biological replicate (*n* = 3) are plotted. P-values from Welch t-test <0.001.

We chose to focus on competition between WT and the overexpression strains since we did not see any significant difference in biofilm or microcolony formation between WT and the deletion mutant in the *in vitro* experiments, and because the lack of fimbriae in *Xenorhabdus* has been shown to not affect colonization ability in previous studies [38]. A small scale experiment with ten replicates where the deletion mutant was competing against WT showed no significant difference in colonization efficiency by CFU counting between the two strains (Figure S6).

To enable quantification of the colonization efficiency of WT versus the *ymdA* overexpression strain, we chromosomally integrated RFP and GFP in both strain backgrounds to create four different strains to test WT-GFP versus *ymdA* overexpression-RFP, and WT-RFP versus *ymdA* overexpression-GFP. By doing this any impact of different burden from expressing the different fluorescent proteins is eliminated. *ymdA* overexpression strains were significantly inhibited in their ability to colonize the nematode compared to WT, as indicated by both microscopy of whole animals (Figure 4B) and CFU counts of ground up nematodes (Figure 4C) where WT bacteria on average colonize more than 80% of the colonized nematodes.

These results show that high levels of *ymdA* negatively impacts the ability of *X. griffiniae* to colonize *Steinernema hermaphroditum*. The downregulation of *ymdA* in *X. griffiniae* when co-cultured with *Steinernema hermaphroditum* might therefore be a response made to actively facilitate successful colonization of the nematode.

## Discussion

A deeper understanding of how bacteria sense nematodes can fill in knowledge gaps regarding signaling and interactions in different ecosystems. Here, we discovered that symbiotic bacteria can sense and respond to the presence of their host nematode in a contact-independent manner. Then, we followed up on the most differentially expressed gene, *ymdA*, coding for a fimbrial protein, that is down-regulated in the presence of nematodes. Our findings provide insight into how symbiotic bacteria respond to host-derived cues, expanding our understanding of host-microbe recognition principles.

*ymdA* had opposing impacts on cell-cell interactions at different scales: at the macroscale, increased *ymdA* inhibited biofilm formation, while at the microscale, increased *ymdA* promoted microcolony clustering and size. One important difference between the two experiments testing biofilm formation is the oxygen concentration, which may play a role in the adherence ability, as seen in previous studies [39, 40]. There is also a possibility that cis-interactions with other fimbriae reduce trans-interactions with surfaces and fimbriae on other bacteria (Figure 5). A likely possibility is that there is an optimal intermediate level of fimbriae that maximize robust cell-cell adherence, where cell refers both to other bacterial cells, but also epithelial cells in the nematode receptacle. Nonetheless, bacteria overexpressing *ymdA* had a colonization defect when competed with WT bacteria, despite having the same colonization efficiency as WT under a non-competitive condition (Figure S7).

**Figure 5:**
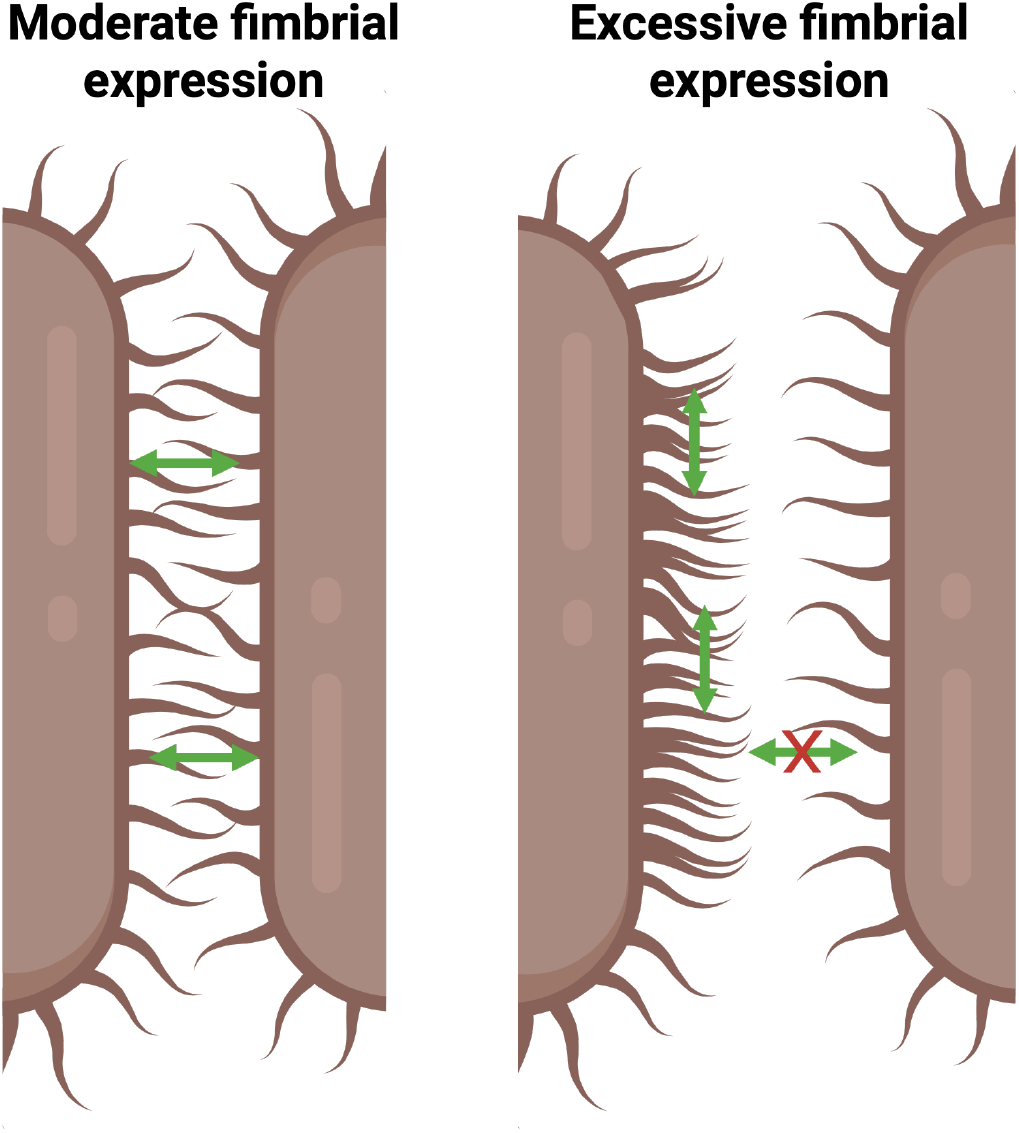
Having too many fimbriae on the cell surface might lead to cis-interactions, preventing trans-interactions that promote cells adhering to each other. Our results suggest that the establishment of the host-symbiont pairing between *Steinernema hermaphroditum* and *X. griffiniae* is not purely a one-sided interaction where the nematode senses and seeks out its symbiont, but might instead be a two-way interaction where both partners sense and respond to the presence of the other in a way that promotes successful colonization and establishment of the symbiosis. However, the molecular mechanism underlying the ability of *X. griffiniae* to sense its host still needs to be elucidated. The fact that *X. griffiniae* only exhibited a transcriptional response to axenic nematodes indicates that the nematode’s secretome may differ between axenic and colonized individuals, as seen in previous studies [41]. These results further suggest the possibility that axenic *S. hermaphroditum* may even actively release chemical cues to signal to *X. griffiniae* that a colonizable host is nearby. Future work to elucidate what metabolites differ in the secretome between colonized and axenic nematodes in this species are needed to fully understand the underlying sensing mechanism used by the bacteria.

Furthermore, understanding nematode sensing in bacteria is the first step in enabling several biotechnology applications. One challenge in agriculture is the widespread infections by root-knot nematodes that cause crop yield losses worth over $80 billion per year [42]. In addition to current methods for detection such as soil collection followed by direct counts or qPCR [43], the construction of whole-cell biosensors for certain species could become an important tool for early detection of outbreaks. Another challenge is the infections caused by insect pests. *Steinernema* and *Heterorhabditis* along with their bacterial symbionts have been suggested as potential replacements or supplements to chemical pesticides in agriculture due to their ability to kill insect larvae [44, 45]. Even though it has been known for decades that these organisms have this capability, we are not near widespread use of them in agriculture. This can be due to regulation, scale-up difficulties etc., but could also be due to them not being efficient enough for this application. Deeper understanding into the establishment of the symbiotic relationship and their insect virulence, might allow us to engineer them for improved efficiency.

## Author contributions

EML and DKN conceived the project idea; EML, MC and DKN designed the experiments; EML, CRM and MC performed the experiments; EML analyzed and visualized the data, and wrote the manuscript; EML, RMM, MC and DKN edited the manuscript.

## Acknowledgements

We thank Dr. Jenny Heppert and Prof. Heidi Goodrich-Blair for sending us the donor vectors used in this study. We thank Prof. Richard Murray for mentorship and valuable feedback throughout this study. We thank Dr. Georgia Squyres for helpful suggestions about image analysis. We thank Dr. Nate Glasser for input on the RNA-sequencing analysis. We thank Prof. Ned Ruby for helpful discussions on biofilm formation and competition. All figures were created using BioRender.com. This work was supported by Caltech’s Resnick Sustainability Institute.

## Conflict of interest

The authors declare no competing financial interest.

## Supplemental Information

Supplemental information document containing:

- List of all strains/plasmids used in this work.
- Volcano plots for conventional WT nematodes and axenic *daf-22* nematodes. -List of genes of differentially regulated genes in the axenic *daf-22* nematode condition.
- More microscopy images from the attachment experiment.
- Box plot from CFU counts of small-scale WT vs deletion mutant colonization experiment.
- Box plot comparing small-scale non-competitive colonization by WT and the overexpression strain.
- Flow cytometry data from quantification of growth on liver kidney plates.
- Nucleotide sequences of hypothetical proteins in the RNA-sequencing experiment

## Notes

https://www.ncbi.nlm.nih.gov/geo/

### Competing Interest Statement

The authors have declared no competing interest.

## References

[1] M. T. Gebremikael, H. Steel, D. Buchan, W. Bert, and S. De Neve, “Nematodes enhance plant growth and nutrient uptake under c and n-rich conditions,” Scientific Reports, vol. 6, no. 1, p. 32862, 2016.

[2] G. W. Yeates, T. Bongers, R. G. De Goede, D. W. Freckman, and S. Georgieva, “Feeding habits in soil nematode families and genera—an outline for soil ecologists,” Journal of Nematology, vol. 25, no. 3, p. 315, 1993.

[3] E. Kenney and I. Eleftherianos, “Entomopathogenic and plant pathogenic nematodes as opposing forces in agriculture,” International Journal for Parasitology, vol. 46, pp. 13–19, Jan. 2016.

[4] S. Brooker, “Estimating the global distribution and disease burden of intestinal nematode infections: adding up the numbers–a review,” International Journal for Parasitology, vol. 40, no. 10, pp. 1137–1144, 2010.

[5] J. I. Choi, K.-h. Yoon, S. Subbammal Kalichamy, S.-S. Yoon, and J. Il Lee, “A natural odor attraction between lactic acid bacteria and the nematode Caenorhabditis elegans,” The ISME Journal, vol. 10, pp. 558–567, Mar. 2016.

[6] D. Prakash, A. Ms, B. Radhika, R. Venkatesan, S. H. Chalasani, and V. Singh, “1-Undecene from Pseudomonas aeruginosa is an olfactory signal for flightor-fight response in Caenorhabditis elegans,” The EMBO Journal, vol. 40, p. e106938, July 2021.

[7] E. Pradel, Y. Zhang, N. Pujol, T. Matsuyama, C. I. Bargmann, and J. J. Ewbank, “Detection and avoidance of a natural product from the pathogenic bacterium Serratia marcescens by Caenorhabditis elegans,” Proceedings of the National Academy of Sciences, vol. 104, no. 7, pp. 2295–2300, 2007.

[8] P. Manosalva, M. Manohar, S. H. Von Reuss, S. Chen, A. Koch, F. Kaplan, A. Choe, R. J. Micikas, X. Wang, K.-H. Kogel, et al., “Conserved nematode signalling molecules elicit plant defenses and pathogen resistance,” Nature Communications, vol. 6, no. 1, p. 7795, 2015.

[9] Y. Yu, Y. K. Zhang, M. Manohar, A. B. Artyukhin, A. Kumari, F. J. Tenjo-Castano, H. Nguyen, P. Routray, A. Choe, D. F. Klessig, and F. C. Schroeder, “Nematode Signaling Molecules Are Extensively Metabolized by Animals, Plants, and Microorganisms,” ACS Chemical Biology, vol. 16, pp. 1050– 1058, June 2021.

[10] L. Zhang, Y. Wei, Y. Tao, S. Zhao, X. Wei, X. Yin, S. Liu, and Q. Niu, “Molecular mechanism of the smart attack of pathogenic bacteria on nematodes,” Microbial Biotechnology, vol. 13, no. 3, pp. 683–705, 2020.

[11] K. E. Murfin, A. C. Whooley, J. L. Klassen, and H. Goodrich-Blair, “Comparison of Xenorhabdus bovienii bacterial strain genomes reveals diversity in symbiotic functions,” BMC Genomics, vol. 16, p. 889, Dec. 2015.

[12] P. Grewal, M. Matsuura, and V. Converse, “Mechanisms of specificity of association between the nematode Steinernema scapterisci and its symbiotic bacterium,” Parasitology, vol. 114, no. 5, pp. 483–488, 1997.

[13] M. Sicard, N. Le Brun, S. Pages, B. Godelle, N. Boemare, and C. Moulia, “Effect of native Xenorhabdus on the fitness of their Steinernema hosts: contrasting types of interaction,” Parasitology Research, vol. 91, pp. 520–524, 2003.

[14] E. E. Herbert and H. Goodrich-Blair, “Friend and foe: the two faces of Xenorhabdus nematophila,” Nature Reviews Microbiology, vol. 5, no. 8, pp. 634– 646, 2007.

[15] J. G. McMullen, B. F. Peterson, S. Forst, H. G. Blair, and S. P. Stock, “Fitness costs of symbiont switching using entomopathogenic nematodes as a model,” BMC Evolutionary Biology, vol. 17, pp. 1–14, 2017.

[16] D. Park, K. Ciezki, R. Van Der Hoeven, S. Singh, D. Reimer, H. B. Bode, and S. Forst, “Genetic analysis of xenocoumacin antibiotic production in the mutualistic bacterium Xenorhabdus nematophila,” Molecular Microbiology, vol. 73, no. 5, pp. 938–949, 2009.

[17] S. Chaudhary, G. Singh, N. Gupta, C. Ghosh, and J. S. Rathore, “New face in the row of bioactive compounds and toxin-antitoxin modules: Xenorhabdus nematophila,” Journal of Asia-Pacific Entomology, vol. 26, no. 4, p. 102148, 2023.

[18] S. Ahmed and Y. Kim, “Differential immunosuppression by inhibiting pla2 affects virulence of Xenorhabdus hominickii and Photorhabdus temperata temperata,” Journal of Invertebrate Pathology, vol. 157, pp. 136–146, 2018.

[19] C. E. Cowles and H. Goodrich-Blair, “The Xenorhabdus nematophila nilABC genes confer the ability of Xenorhabdus spp. to colonize Steinernema carpocapsae nematodes,” Journal of Bacteriology, vol. 190, no. 12, pp. 4121–4128, 2008.

[20] M. Cao, H. T. Schwartz, C.-H. Tan, and P. W. Sternberg, “The entomopathogenic nematode Steinernema hermaphroditum is a self-fertilizing hermaphrodite and a genetically tractable system for the study of parasitic and mutualistic symbiosis,” Genetics, vol. 220, p. iyab170, Jan. 2022.

[21] G. White, “A method for obtaining infective nematode larvae from cultures,” Science, vol. 66, no. 1709, pp. 302–303, 1927.

[22] T. Stiernagle, “Maintenance of c. elegans,” WormBook: The online review of C. elegans biology [Internet], 2006.

[23] J. K. Heppert, R. M. Awori, M. Cao, G. Chen, J. McLeish, and H. GoodrichBlair, “Analyses of Xenorhabdus griffiniae genomes reveal two distinct subspecies that display intra-species variation due to prophages,” BMC Genomics, vol. 25, no. 1, p. 1087, 2024.

[24] T. Seemann, “Prokka: rapid prokaryotic genome annotation,” Bioinformatics, vol. 30, no. 14, pp. 2068–2069, 2014.

[25] N. L. Bray, H. Pimentel, P. Melsted, and L. Pachter, “Near-optimal probabilistic RNA-seq quantification,” Nature Biotechnology, vol. 34, no. 5, pp. 525–527, 2016.

[26] M. I. Love, W. Huber, and S. Anders, “Moderated estimation of fold change and dispersion for RNA-seq data with DESeq2,” Genome Biology, vol. 15, pp. 1–21, 2014.

[27] A. D. Halleran, A. Swaminathan, and R. M. Murray, “Single day construction of multigene circuits with 3G assembly,” ACS Synthetic Biology, vol. 7, no. 5, pp. 1477– 1480, 2018.

[28] G. A. O’Toole, “Microtiter dish biofilm formation assay,” Journal of Visualized Experiments: JoVE, no. 47, p. 2437, 2011.

[29] N. M. St Thomas, T. G. Myers, O. S. Alani, H. Goodrich-Blair, and J. K. Heppert, “Green and red fluorescent strains of Xenorhabdus griffiniae HGB2511, the bacterial symbiont of the nematode Steinernema hermaphroditum (india),” microPublication Biology, vol. 2024, pp. 10–17912, 2024.

[30] K. E. Murfin, A. R. Dillman, J. M. Foster, S. Bulgheresi, B. E. Slatko, P. W. Sternberg, and H. Goodrich-Blair, “Nematode-Bacterium Symbioses— Cooperation and Conflict Revealed in the “Omics” Age,” The Biological Bulletin, vol. 223, pp. 85–102, Aug. 2012.

[31] M. Cao, “CRISPR-Cas9 genome editing in Steinernema entomopathogenic nematodes,” bioRxiv, pp. 2023–11, 2023.

[32] A. Renda, S. Poly, Y.-J. Lai, A. Pannuri, H. Yakhnin, A. H. Potts, P. C. Bevilacqua, T. Romeo, and P. Babitzke, “CsrA-Mediated Translational Activation of ymdA Expression in Escherichia coli,” mBio, vol. 11, pp. e00849–20, Oct. 2020.

[33] M. Kim and K.-s. Kim, “Stress-responsively modulated ymdAB-clsC operon plays a role in biofilm formation and apramycin susceptibility in Escherichia coli,” FEMS Microbiology Letters, vol. 364, July 2017.

[34] R. B. McLay, H. N. Nguyen, Y. A. Jaimes-Lizcano, N. K. Dewangan, S. Alexandrova, D. F. Rodrigues, P. C. Cirino, and J. C. Conrad, “Level of fimbriation alters the adhesion of Escherichia coli bacteria to interfaces,” Langmuir, vol. 34, no. 3, pp. 1133–1142, 2018.

[35] Q. Liu, J. Zhu, N. Liu, W. Sun, B. Yu, H. Niu, D. Liu, P. Ouyang, H. Ying, Y. Chen, et al., “Type i fimbriae subunit fima enhances Escherichia coli biofilm formation but affects l-threonine carbon distribution,” Frontiers in Bioengineering and Biotechnology, vol. 10, p. 904636, 2022.

[36] C.-L. Y. Ong, G. C. Ulett, A. N. Mabbett, S. A. Beatson, R. I. Webb, W. Monaghan, G. R. Nimmo, D. F. Looke, A. G. McEwan, and M. A. Schembri, “Identification of type 3 fimbriae in uropathogenic Escherichia coli reveals a role in biofilm formation,” Journal of Bacteriology, vol. 190, no. 3, pp. 1054–1063, 2008.

[37] M. A. Lasaro, N. Salinger, J. Zhang, Y. Wang, Z. Zhong, M. Goulian, and J. Zhu, “F1C fimbriae play an important role in biofilm formation and intestinal colonization by the Escherichia coli commensal strain Nissle 1917,” Applied and Environmental Microbiology, vol. 75, no. 1, pp. 246–251, 2009.

[38] H. Snyder, H. He, H. Owen, C. Hanna, and S. Forst, “Role of Mrx Fimbriae of Xenorhabdus nematophila in Competitive Colonization of the Nematode Host,” Applied and Environmental Microbiology, vol. 77, pp. 7247–7254, Oct. 2011.

[39] S.-J. Ahn and R. A. Burne, “Effects of oxygen on biofilm formation and the atla autolysin of Streptococcus mutans,” Journal of Bacteriology, vol. 189, no. 17, pp. 6293–6302, 2007.

[40] J. J. Cotter, J. P. O’Gara, D. Mack, and E. Casey, “Oxygen-mediated regulation of biofilm development is controlled by the alternative sigma factor σb in staphylococcus epidermidis,” Applied and environmental microbiology, vol. 75, no. 1, pp. 261–264, 2009.

[41] E. Lefoulon, J. G. McMullen, and S. P. Stock, “Transcriptomic analysis of Steinernema nematodes highlights metabolic costs associated to Xenorhabdus endosymbiont association and rearing conditions,” Frontiers in Physiology, vol. 13, p. 821845, 2022.

[42] G. C. Bernard, M. Egnin, and C. Bonsi, “The impact of plant-parasitic nematodes on agriculture and methods of control,” in Nematology-concepts, Diagnosis and Control, IntechOpen, 2017.

[43] A. Braun-Kiewnick, N. Viaene, L. Folcher, F. Ollivier, G. Anthoine, B. Niere, M. Sapp, B. van de Vossenberg, H. Toktay, and S. Kiewnick, “Assessment of a new qPCR tool for the detection and identification of the root-knot nematode Meloidogyne enterolobii by an international test performance study,” European Journal of Plant Pathology, vol. 144, pp. 97–108, 2016.

[44] I. Kepenekci, S. Hazir, E. Oksal, and E. E. Lewis, “Application methods of Steinernema feltiae, Xenorhabdus bovienii and Purpureocillium lilacinum to control root-knot nematodes in greenhouse tomato systems,” Crop Protection, vol. 108, pp. 31–38, 2018.

[45] S. Manochaya, S. Udikeri, B. S. Srinath, M. Sairam, S. V. Bandlamori, and K. Ramakrishna, “In vivo culturing of entomopathogenic nematodes for biological control of insect pests: A review,” Journal of Natural Pesticide Research, vol. 1, p. 100005, 2022.

