## Supplemental Tables and Figures for "The nematode symbiotic bacterium *Xenorhabdus griffiniae* can sense and respond to the presence of its host *Steinernema hermaphroditum*"

**Table S1**: Plasmids used in this study.

| Plasmid | Antibiotic marker | Description | Source |
| --- | --- | --- | --- |
| pRK100 | CmR | Ori6K suicide vector for gene deletions | Provided by Heidi Goodrich-Blair lab |
| pUX-B13 | AmpR | Helper plasmid for Tn7 insertion | (Bao et al. 1991) |
| HGB1262 | KanR, AmpR, CmR, SmR | Tn7 donor plasmid with GFPmut3 | (St Thomas et al. 2024) |
| EML002 | KanR, AmpR | Derivative of HGB1262 | (Larsson, Wang, and Murray 2025) |
| EML006 | SmR, AmpR | Derivative of HGB1262 with removed KanR cassette - GFPmut3 donor | (Larsson, Wang, and Murray 2025) |
| EML007 | SmR, AmpR | Derivative of HGB1262 with removed KanR cassette - TurboRFP donor | (Larsson, Wang, and Murray 2025) |
| EML047 | KanR, CmR | Donor for ymdA1/2 knockout | This study |
| EML048 | KanR, AmpR | Donor for ymdA1/2 overexpression strain | This study |
| EML008 | KanR, AmpR | Donor for GFPmut3 | This study |
| EML011 | KanR, AmpR | Donor for TurboRFP | This study |

CmR: chloramphenicol resistance cassette, KanR: kanamycin resistance cassette, AmpR: ampicillin resistance cassette, SmR: streptomycin resistance cassette

**Table S2**: Strains used in this study.

| Strain | Description | Source |
| --- | --- | --- |
| HGB2511 | *Xenorhabdus griffiniae* wildtype strain isolated from *Steinernema hermaphroditum* | (Cao et al. 2022) |
| GYC53 | *E. coli* MFDpir (Ferrières et al. 2010)(Ferrières et al. 2010) with helper plasmid | Provided by Margaret McFall-Ngai lab |

| | | |
|---|---|---|
| | pUX-B13 | |
| HGB1262 | *E. coli* conjugal donor strain | (St Thomas et al. 2024) |
| xEML047 | *Xenorhabdus griffiniae ymdA* deletion strain | This study |
| xEML016 | *Xenorhabdus griffiniae ymdA* overexpression strain | This study |
| xEML048 | *Xenorhabdus griffiniae ymdA* deletion strain with GFP marker | This study |
| xEML049 | *Xenorhabdus griffiniae ymdA* deletion strain with RFP marker | This study |
| xEML032 | *Xenorhabdus griffiniae* wildtype strain with GFP marker | This study |
| xEML013 | *Xenorhabdus griffiniae* wildtype strain with RFP marker | This study |
| xEML035 | *Xenorhabdus griffiniae ymdA* overexpression strain with GFP marker | This study |
| xEML034 | *Xenorhabdus griffiniae ymdA* overexpression strain with RFP marker | This study |

**Table S3**: Genes differentially expressed for bacteria grown +/- axenic *daf-22* nematodes

| **Gene annotation** | **Literature** | **Reference** |
|---|---|---|
| *ymdA* | Type 1 fimbrial protein | (Kim and Kim 2017; Renda et al. 2020) |
| *ybhF* | Multidrug ABC transporter ATP-binding protein | (Yamanaka et al. 2016; Feng et al. 2020) |
| Hypothetical protein 343 | BLASTx: 100% identity with hlyD secretion protein | NCBI: WP_189758156.1 |
| *cecR* | Regulator for chloramphenicol sensitivity in *E. coli* | (Yamanaka et al. 2016) |

| | | |
|---|---|---|
| *dps* | DNA protection | (Nair and Finkel 2004) |
| *moeZ* | Molybdopterin-synthase adenylyltransferase | |
| *cheW* | Chemotaxis | (Liu and Parkinson 1989) |
| *glnA* | Glutamine synthase | (Woolfolk, Shapiro, and Stadtman 1966) |
| *tag* | DNA-3-methyladenine glycosylase 1 | |
| *amtB* | Ammonium transporter | (Soupene et al. 1998) |
| *motA* | Motility | (Dean et al. 1984) |

**Table S4**: IJ density and viability in the trans-well experiment.

| IJ | Number of IJs in transwell | Viability before experiment | Viability after experiment |
|---|---|---|---|
| conventional WT-1 | 2041 | 1 | 0.995 |
| conventional WT-2 | 3375 | 0.995 | 0.99 |
| conventional WT-3 | 4541 | 0.99 | 0.902 |
| Axenic IJ-1 | 7250 | 0.995 | 0.975 |
| Axenic IJ-2 | 6333 | 0.99 | 0.957 |
| Axenic IJ-3 | 5250 | 0.99 | 0.956 |
| Axenic *daf-22*-1 | 6042 | 0.995 | 0.949 |
| Axenic *daf-22-2* | 3708 | 0.985 | 0.975 |
| Axenic *daf-22-3* | 5250 | 0.995 | 0.97 |

**Figure S1: Volcano plot for bacteria grown +/- WT colonized nematodes**

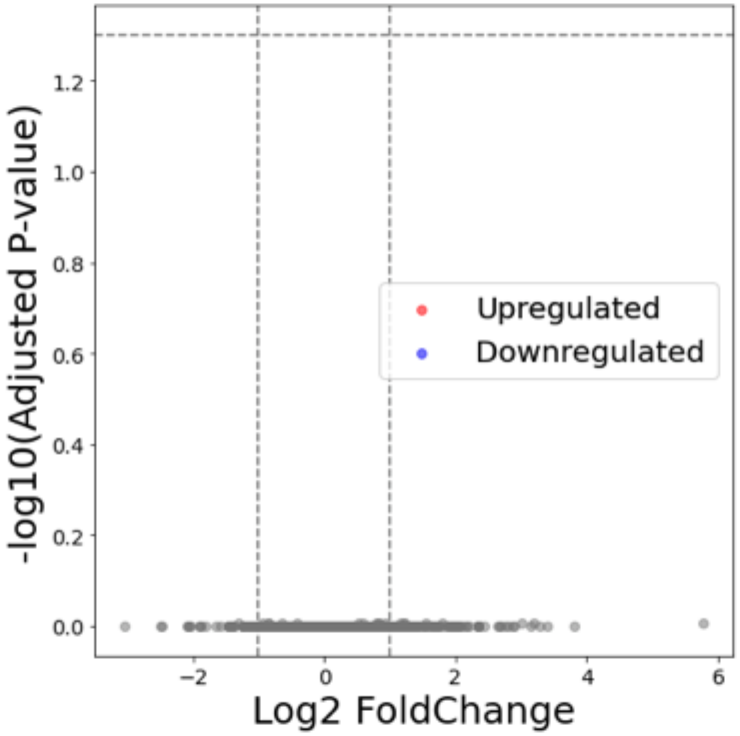


*Volcano plot generated from RNA-sequencing data for three biological replicates of bacteria exposed to conventional (colonized) nematodes.*

**Figure S2: Volcano plot for bacteria grown +/- axenic *daf-22* deficient nematodes**

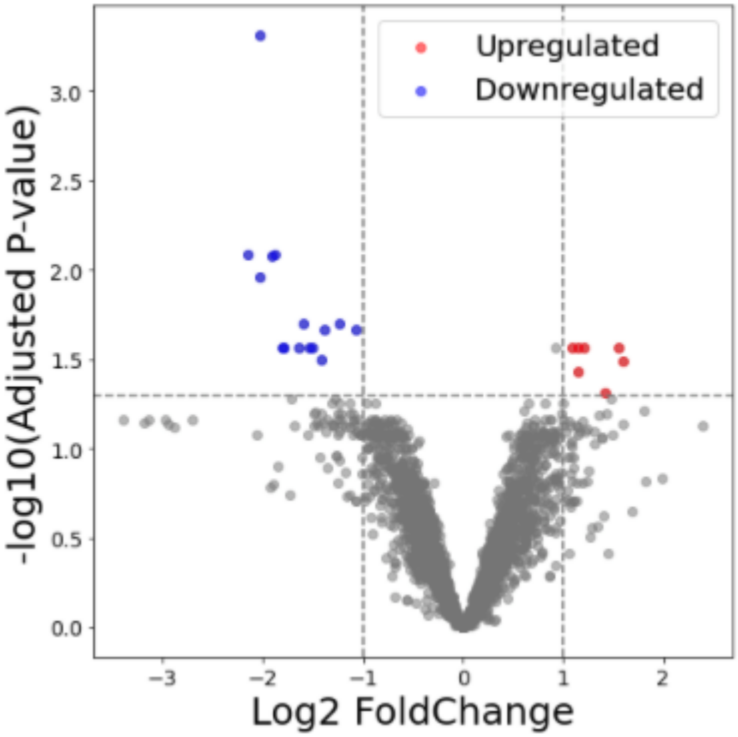


*Volcano plot generated from RNA-sequencing data for three biological replicates of bacteria exposed to axenic daf-22 mutant nematodes.*

**Figure S3: Additional images from the micro colony experiment in Figure 3**

**WT**

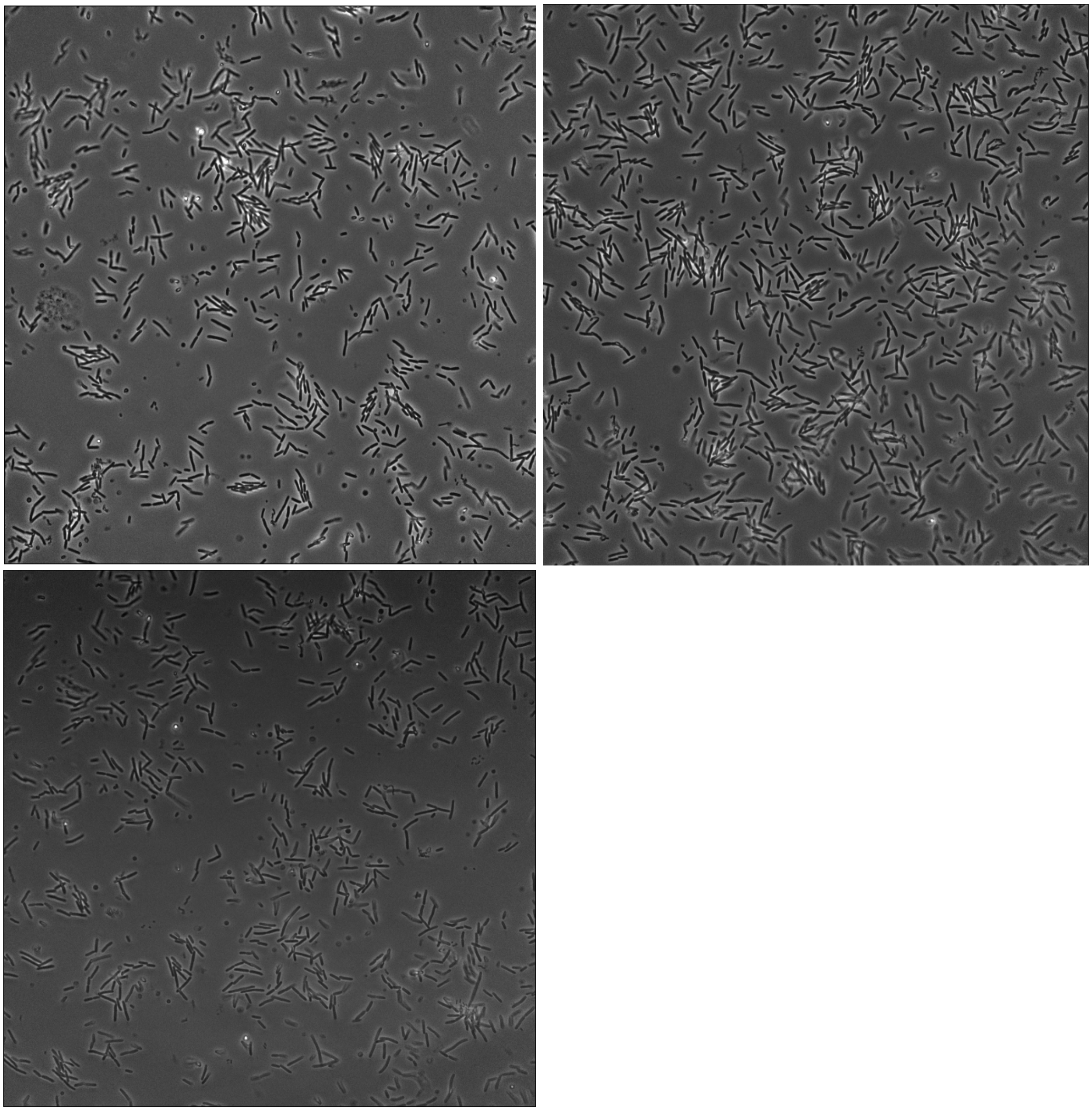

**Deletion mutant**

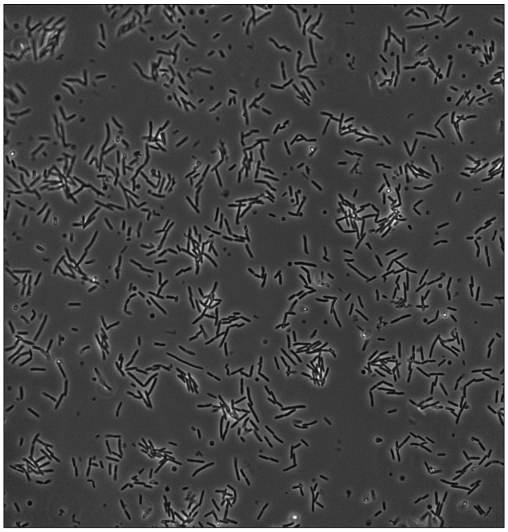

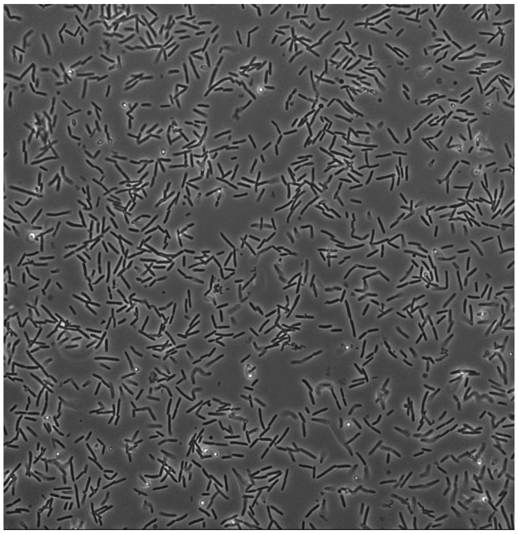

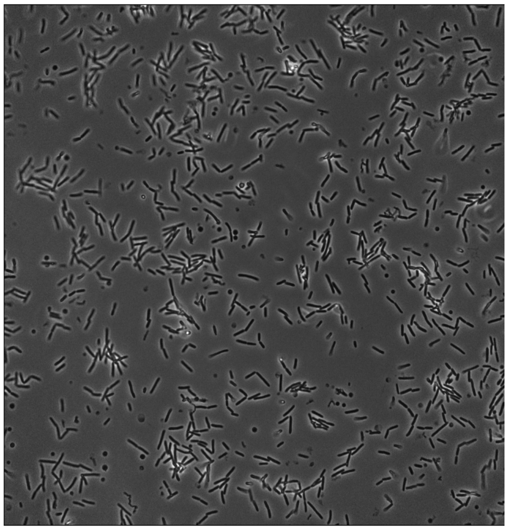

***ymdA* overexpression strain**

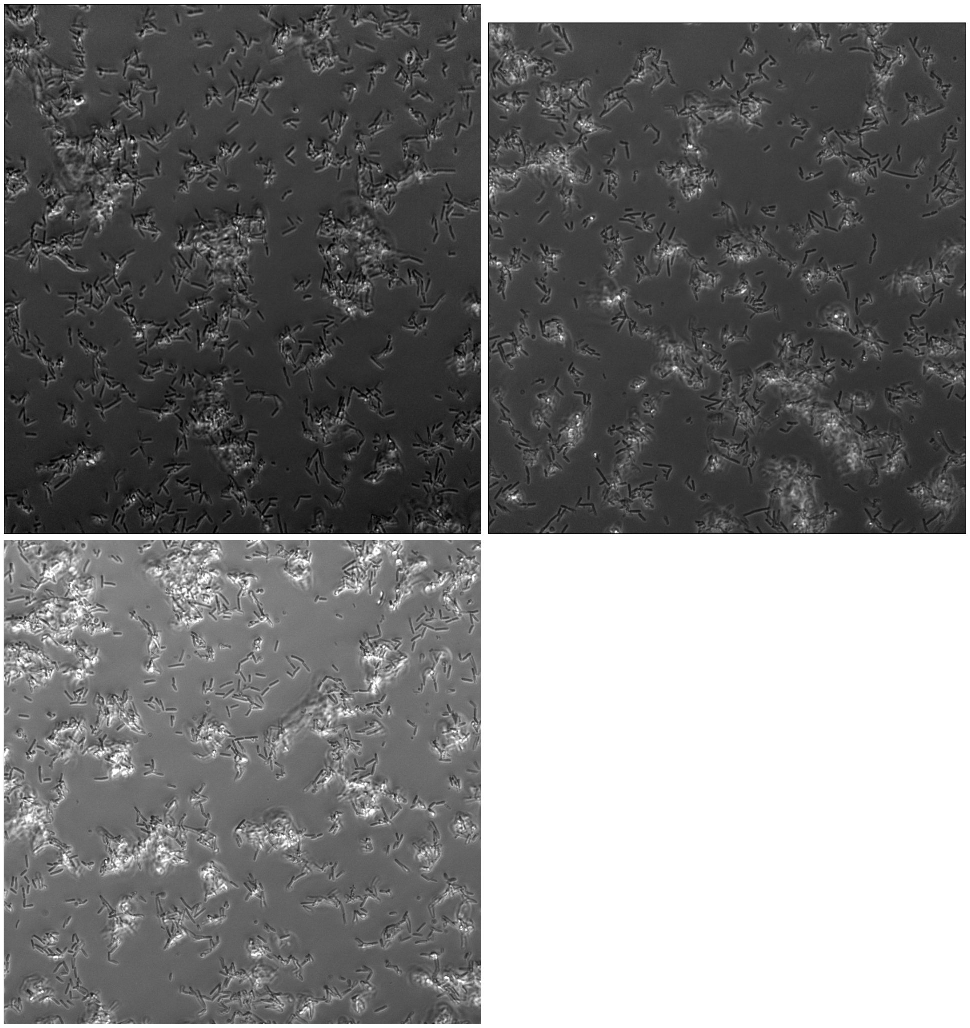

**Figure S5: Flow cytometry control quantification of bacteria grown on liver-kidney plates**

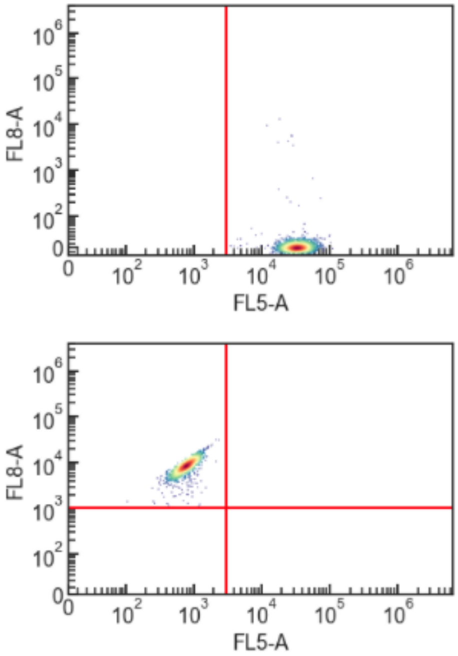


FL-8 (RFP) and FL-5 (GFP) flow cytometry measurements of bacteria scraped off from a liver-kidney plate after incubation. Here wildtype *Xenorhabdus griffiniae* are marked with RFP and the *ymdA* overexpression strain is marked with GFP. The number of cells counted in the RFP channel: 1980, the number of cells counted in the GFP channel: 2234. This is indicating similar growth rates when co-cultured on plates.

**Figure S6: Colonization when WT and the deletion strain colonize axenic nematodes competitively by CFU counts**

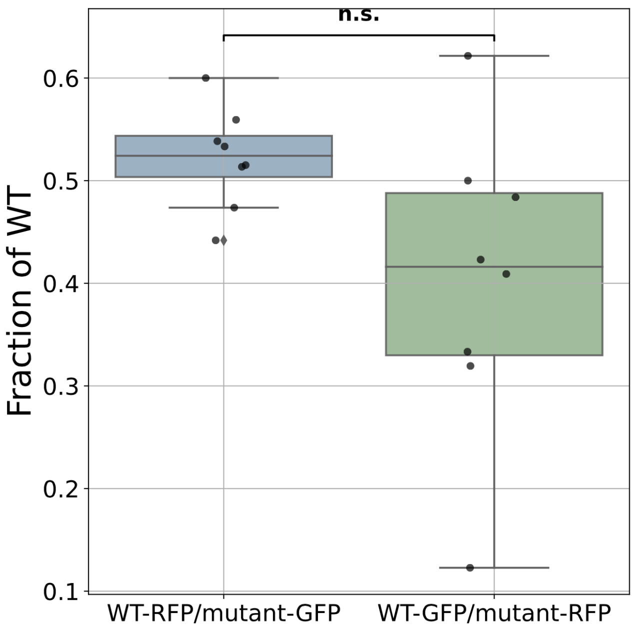


*Four replicates per pairing plotted.*

**Figure S7: Colonization frequency when WT and the overexpression strain colonize axenic nematodes without competition by whole nematode counts**

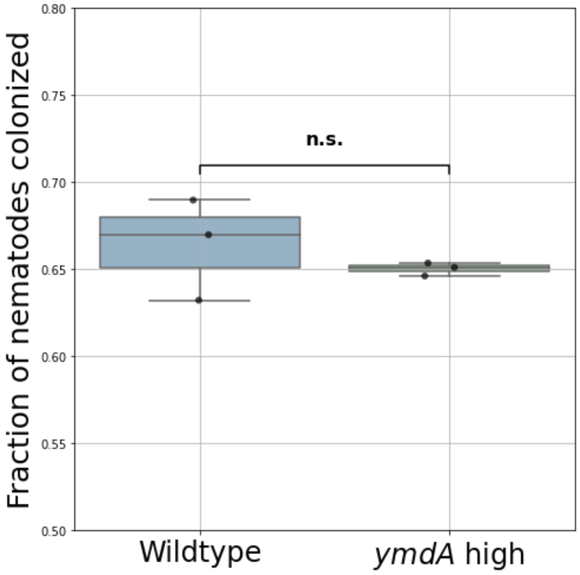


*3 replicates per strain plotted.*

### F: Nucleotide sequences of hypothetical proteins

#### Hypothetical protein 280

ATGATAAAGCTACGGCGGGTTCTGATTGTCGGTTCAAAATTTGGTGAACTGTATCTTAATGCGTTTCTACAGCCTCGGTCA
GATCTGGAACTGGCCGGAATTCTGGCGCAAGGAAGCCCTCGTTCACAACAACTTGCCCGCGATTTCAATGTCCCCCTATT
CCGTTCATTGGATGAATTGCCGGATGATATCGATATTGCCTGTGTGGTCATACGCGCAGAGGTTATTGGTGGCGAAGGCG
ACCAATTAACTCAGGCGTTATTGGCACGGGGCATTTCGGTCATTCAGGAACATCCTCTTGATGCAGAAGCGATCAAAAGA
CACCAATCAACAGCTAACCAGCATAATGCTATGTTTTGGGTAAACAGCTTTTACAGCCAATGTCAGGCTGGCAGAAGTTG
GTTACATGCGGCTCATCAAATCGAGCAATTATCACAGCAACGTCCGGTAAGTGGCTATCTGGCAACCAGCCGCCAGTTAC
TTTTTTCCGCCCTTGATCTGTTGTTACAGGGCTGTCATTGCCCGGAACAGATTGAGATAGCTTATGTAGGTCAGCGAAACA
ATGCATTTCATCGCTTTGAACTCAGCATAGCGGGAACCACAATAACGTTAGATGTTCAATCCTATCTGGACCCCAGGGAT
CCCGATATGCACAACCTGGTCATGCATAAAATGACGCTAATTTGGCCGGCCGGTTATCTGACGCAGGAAGCCAGCTATGG
ACCGGTTATCTGGACACCGGTTCTTCATGCACCAAACCATCTGAATAATGAACAAAGTCTTTACCTGAATGCCGGCCAGC
CGGAAGGCGCTTATCTGCACCAGACAACATCACAAATTCTTTGTCCGGCAACGCAAAATTGGATGACTGCATTTGAATGT
GATGGTGCCGAAGGTGTGGCACGCCTCCTCAATACCTTGAATTCAGTATTGAATGGGGCACTCTGCCCCCCTGCTTTTAG
CTACGATTACCAATTGCGGCTGGCAGGCTTGTGGCAACAAGTTTTACGCATGATGGGAGAGCCGGTTACTCAAGAGCTAC
AGCCACCCGTCTGGATTGATCCAACCCAATTGATGTTAATTAAGGAAGAAAACTGA

#### Hypothetical protein 343

ATGAACAGTAAAAAGTTTGCTCTTGCCCTATTGGCACTCATTGTTGTTATCAGTGCTATTGTCGGTATCTACTATTATCAG
GAACAAAATAGCAGGGAGCTTACCCTGTATGGCAATGTTGATGTCAGGACTGTTAACCTAAGTTTTCGGGTTGCAGGTAA
GCTGGCTAATTTGCAGGTGGATGAAGGTGATGCCATTACTGTGGGCCAAATGATTGGTCGCCTTGATGATGCCCCCTTTA
TCAATGCCTTGAATCAAGCCAAAGCAGCTCGTGATGTCGCTAAAGCAAATCTTGCCAAAGCCGAAGCGGGTTATCGCAGC
GAAGAAATCGCCCAGGTACGTGCTGAAGTCAAGCAAAAAGAGTCAGCATTCCATTTTGCCGATAGTTTCCTGAAACGCCA
GCAAGGGTTATGGCAGAGCAAAACAATCTCAGCCAATGATCTGGAAAATGCCAGAACCGGACGCAATCAAGCACTGGCC
GCTTACCAGGCAGCAAAAGATAAATTGCGTCAGTTCGAAACTGGTTATCGTATAGAAGAAATTGCAGCGGCAAAAGCCCA
ACTTATTCAAGCAGAAGCAGCCGTGGCACAGGCTGAACTTAATCTAAAAGATACTCAACTCACTTCTCCCTCGGCAGGGA
TGATCCTGACCAGAGCCATTGAGCCAGGCACCATGCTGGCGGCTGGCAACACGGTTTTTACCCTGTCATTGACCAATCCT
GTTTGGATCAGGGCATATATTGATGAGCCTAATCTTAATCAGGCTATCGCTGGACGGGAGGTCTTGATTTATACCGATGG
GCGGAAAAACCAGCCTTATCATGGCAAAATCGGTTTCGTTTCTCCAACTGCCGAATTCACCCCAAAAAATGTCGAAACCC
CTGACTTAAGAACAGACTTGGTCTATCGCCTGCGCATTATTGTCCTTGATCCCGATGAAGGACTGAAACAAGGAATGCCT
GTTACTTTGCGTTTTACTGAAACAAATAAGTAG

#### Hypothetical protein 588

ATGATACTTGAATCTAAATTTTTTAGCATTTTGAGCCCAGATATAACGGGACCATTGACGCGTAGTGCGTCAAATTTAATA
AATACGCCTTCCAGGGGATTTAATATTAAGAATTTTATCACTATCAGAGAGGTTGAGCTTCAAGAGGCTTTATGTGTTGCT
AATCAAATAAGAACGAACATGATTAATGCTGATTGGTTGGTAAGTGCTAATCAGAACAGACAATTAAGTCAGGAGCAGGA
ACGATGGAATTCGAGGTATAAGGCTAGCTATAGAATACTTTCCCAAATCTCTAGTCTCTTAGAAAGTAACTTTGCAGGTGT
TTCAGCTTCAATAGAACATTATTCCTTTGTTATGTTTTTTCAAGGAAAACCAATTGGTATGTTGATATTTTCTAATAACAAG
GAAAAAGTCACAGAACCTTCTTCTATTATAGAGTTTGCTACCCATATAGGTATTAGAAATAGTGGAATTTTTCTAATGGAA
TATGCGGTAAACAAATCAAAAACACTGGGTAAAAATGGGAATATCAAACTAATACCAGCGCCTGCTGCGAGAAATGTTTA
CTTTCAATATGGATTTAGATATCAAGGAGGTTACATGGTATTAGAACCAGACAAAAGTGATAAATGGGGATTTTGGCGGG
GAGAATATTATTTTAAGGCAAATTGTGTATAA

### References


Bao, Y., D. P. Lies, H. Fu, and G. P. Roberts. 1991. “An Improved Tn7-Based System for the Single-Copy Insertion of Cloned Genes into Chromosomes of Gram-Negative Bacteria.” *Gene* 109 (1): 167–68.

Cao, Mengyi, Hillel T. Schwartz, Chieh-Hsiang Tan, and Paul W. Sternberg. 2022. “The Entomopathogenic Nematode Steinernema Hermaphroditum Is a Self-Fertilizing Hermaphrodite and a Genetically Tractable System for the Study of Parasitic and Mutualistic Symbiosis.” *Genetics* 220 (1). https://doi.org/10.1093/genetics/iyab170.

Dean, G. E., R. M. Macnab, J. Stader, P. Matsumura, and C. Burks. 1984. “Gene Sequence and Predicted Amino Acid Sequence of the motA Protein, a Membrane-Associated Protein Required for Flagellar Rotation in Escherichia Coli.” *Journal of Bacteriology* 159 (3): 991–99.

Feng, Zhenyue, Defu Liu, Lizi Wang, Yanhong Wang, Zhongjing Zang, Zhenhua Liu, Baifen Song, et al. 2020. “A Putative Efflux Transporter of the ABC Family, YbhFSR, in Functions in Tetracycline Efflux and Na(Li)/H Transport.” *Frontiers in Microbiology* 11 (April):556.

Ferrières, Lionel, Gaëlle Hémery, Toan Nham, Anne-Marie Guérout, Didier Mazel, Christophe Beloin, and Jean-Marc Ghigo. 2010. “Silent Mischief: Bacteriophage Mu Insertions Contaminate Products of Escherichia Coli Random Mutagenesis Performed Using Suicidal Transposon Delivery Plasmids Mobilized by Broad-Host-Range RP4 Conjugative Machinery.” *Journal of Bacteriology* 192 (24): 6418–27.

Kim, Moonjeong, and Kwang-Sun Kim. 2017. “Stress-Responsively Modulated ymdAB-clsC Operon Plays a Role in Biofilm Formation and Apramycin Susceptibility in Escherichia Coli.” *FEMS Microbiology Letters* 364 (13). https://doi.org/10.1093/femsle/fnx114.

Larsson, Elin M., Olivia Y. Wang, and Richard M. Murray. 2025. “A DNA Part Library for Reliable Engineering of the Emerging Model Nematode Symbiotic Bacterium Xenorhabdus Griffiniae HGB2511.” *bioRxiv*. https://doi.org/10.1101/2025.06.09.658710.

Liu, J. D., and J. S. Parkinson. 1989. “Role of CheW Protein in Coupling Membrane Receptors to the Intracellular Signaling System of Bacterial Chemotaxis.” *Proceedings of the National Academy of Sciences of the United States of America* 86 (22): 8703–7.

Nair, Sudha, and Steven E. Finkel. 2004. “Dps Protects Cells against Multiple Stresses during Stationary Phase.” *Journal of Bacteriology* 186 (13): 4192–98.

Renda, Andrew, Stephanie Poly, Ying-Jung Lai, Archana Pannuri, Helen Yakhnin, Anastasia H. Potts, Philip C. Bevilacqua, Tony Romeo, and Paul Babitzke. 2020. “CsrA-Mediated Translational Activation of Expression in Escherichia Coli.” *mBio* 11 (5). https://doi.org/10.1128/mBio.00849-20.

Soupene, E., L. He, D. Yan, and S. Kustu. 1998. “Ammonia Acquisition in Enteric Bacteria: Physiological Role of the Ammonium/methylammonium Transport B (AmtB) Protein.” *Proceedings of the National Academy of Sciences of the United States of America* 95 (12): 7030–34.

St Thomas, Nadia M., Tyler G. Myers, Omar S. Alani, Heidi Goodrich-Blair, and Jennifer K. Heppert. 2024. “Green and Red Fluorescent Strains of HGB2511, the Bacterial Symbiont of the Nematode (India).” *microPublication Biology* 2024 (February). https://doi.org/10.17912/micropub.biology.001064.

Woolfolk, C. A., B. Shapiro, and E. R. Stadtman. 1966. “Regulation of Glutamine Synthetase. I. Purification and Properties of Glutamine Synthetase from Escherichia Coli.” *Archives of Biochemistry and Biophysics* 116 (1): 177–92.

Yamanaka, Yuki, Tomohiro Shimada, Kaneyoshi Yamamoto, and Akira Ishihama. 2016. “Transcription Factor CecR (YbiH) Regulates a Set of Genes Affecting the Sensitivity of Escherichia Coli against Cefoperazone and Chloramphenicol.” *Microbiology (Reading, England)* 162 (7): 1253–64.

Bao, Y., D. P. Lies, H. Fu, and G. P. Roberts. 1991. “An Improved Tn7-Based System for the Single-Copy Insertion of Cloned Genes into Chromosomes of Gram-Negative Bacteria.” Gene 109 (1): 167–68.
Cao, Mengyi, Hillel T. Schwartz, Chieh-Hsiang Tan, and Paul W. Sternberg. 2022. “The Entomopathogenic Nematode Steinernema Hermaphroditum Is a Self-Fertilizing Hermaphrodite and a Genetically Tractable System for the Study of Parasitic and Mutualistic Symbiosis.” Genetics 220 (1). https://doi.org/10.1093/genetics/iyab170.
Dean, G. E., R. M. Macnab, J. Stader, P. Matsumura, and C. Burks. 1984. “Gene Sequence and Predicted Amino Acid Sequence of the motA Protein, a Membrane-Associated Protein Required for Flagellar Rotation in Escherichia Coli.” Journal of Bacteriology 159 (3): 991–99.
Feng, Zhenyue, Defu Liu, Lizi Wang, Yanhong Wang, Zhongjing Zang, Zhenhua Liu, Baifen Song, et al. 2020. “A Putative Efflux Transporter of the ABC Family, YbhFSR, in Functions in Tetracycline Efflux and Na(Li)/H Transport.” Frontiers in Microbiology 11 (April):556.
Ferrières, Lionel, Gaëlle Hémery, Toan Nham, Anne-Marie Guérout, Didier Mazel, Christophe Beloin, and Jean-Marc Ghigo. 2010. “Silent Mischief: Bacteriophage Mu Insertions Contaminate Products of Escherichia Coli Random Mutagenesis Performed Using Suicidal Transposon Delivery Plasmids Mobilized by Broad-Host-Range RP4 Conjugative Machinery.” Journal of Bacteriology 192 (24): 6418–27.
Kim, Moonjeong, and Kwang-Sun Kim. 2017. “Stress-Responsively Modulated ymdAB-clsC Operon Plays a Role in Biofilm Formation and Apramycin Susceptibility in Escherichia Coli.” FEMS Microbiology Letters 364 (13). https://doi.org/10.1093/femsle/fnx114.
Larsson, Elin M., Olivia Y. Wang, and Richard M. Murray. 2025. “A DNA Part Library for Reliable Engineering of the Emerging Model Nematode Symbiotic Bacterium Xenorhabdus Griffiniae HGB2511.” bioRxiv. https://doi.org/10.1101/2025.06.09.658710.
Liu, J. D., and J. S. Parkinson. 1989. “Role of CheW Protein in Coupling Membrane Receptors to the Intracellular Signaling System of Bacterial Chemotaxis.” Proceedings of the National Academy of Sciences of the United States of America 86 (22): 8703–7.
Nair, Sudha, and Steven E. Finkel. 2004. “Dps Protects Cells against Multiple Stresses during Stationary Phase.” Journal of Bacteriology 186 (13): 4192–98.
Renda, Andrew, Stephanie Poly, Ying-Jung Lai, Archana Pannuri, Helen Yakhnin, Anastasia H. Potts, Philip C. Bevilacqua, Tony Romeo, and Paul Babitzke. 2020. “CsrA-Mediated Translational Activation of Expression in Escherichia Coli.” mBio 11 (5). https://doi.org/10.1128/mBio.00849-20.
Soupene, E., L. He, D. Yan, and S. Kustu. 1998. “Ammonia Acquisition in Enteric Bacteria: Physiological Role of the Ammonium/methylammonium Transport B (AmtB) Protein.” Proceedings of the National Academy of Sciences of the United States of America 95 (12): 7030–34.
St Thomas, Nadia M., Tyler G. Myers, Omar S. Alani, Heidi Goodrich-Blair, and Jennifer K. Heppert. 2024. “Green and Red Fluorescent Strains of HGB2511, the Bacterial Symbiont of the Nematode (India).” microPublication Biology 2024 (February). https://doi.org/10.17912/micropub.biology.001064.
Woolfolk, C. A., B. Shapiro, and E. R. Stadtman. 1966. “Regulation of Glutamine Synthetase. I. Purification and Properties of Glutamine Synthetase from Escherichia Coli.” Archives of Biochemistry and Biophysics 116 (1): 177–92.
Yamanaka, Yuki, Tomohiro Shimada, Kaneyoshi Yamamoto, and Akira Ishihama. 2016. “Transcription Factor CecR (YbiH) Regulates a Set of Genes Affecting the Sensitivity of Escherichia Coli against Cefoperazone and Chloramphenicol.” Microbiology (Reading, England) 162 (7): 1253–64.
